# Endometrial adhesion G protein-coupled receptors are dynamically expressed across the menstrual cycle and expression is altered by ovarian stimulation

**DOI:** 10.1101/2022.12.04.519044

**Authors:** Nischelle Kalakota, Alexander Lemenze, Lea George, Qingshi Zhao, Tracy Wu, Sara S. Morelli, Nataki C. Douglas, Andy V. Babwah

**Author notes:** Co-senior authors with equal contributions (Nataki C. Douglas, Andy V. Babwah). **Corresponding author**: Dr. Nischelle Kalakota, Department of Obstetrics, Gynecology & Reproductive Health, Rutgers New Jersey Medical School, 185 South Orange Avenue, MSB, E-506, Newark, NJ 07101.

## Abstract

Ovarian stimulation (OS), utilized for the development of multiple ovarian follicles for IVF, induces supraphysiologic levels of E2 and an early rise in P4 that disrupt endometrial differentiation and decreases implantation rates or result in placental insufficiency and pregnancy complications. To improve pregnancy rates and reduce the risk of pregnancy complications associated with IVF, it is crucial to advance our molecular understanding of the molecular regulation of endometrial differentiation. Previous studies from our laboratory suggest G protein-coupled receptors (GPCRs) are important regulators of endometrial differentiation. To investigate this further, using a retrospective dataset, we identified all GPCRs expressed across the proliferative and secretory phase of the menstrual cycle and found that many members of the adhesion G protein-coupled receptor (ADGR) family are dynamically expressed. For each ADGR subfamily exhibiting differentially-expressed genes across the cycle, their expression was investigated by RT-PCR in the non-pregnant mouse uterus and decidua on E7.5 of pregnancy. For those genes expressed in the E7.5 decidua, their expression was further quantified by qPCR across early mouse pregnancy. The RT-PCR screen revealed expression of 13 ADGRs (4 of the 9 subfamilies) in E7.5 decidua and among these genes, many were differentially expressed between E0.5 and E5.5 or 6.5 and between E5.5 and E6.5. The dynamic expression of the ADGRs across the menstrual cycle and in early mouse pregnancy, suggests these *ADGRs* are E2- and/or P4-regulated genes. We therefore hypothesized that for these *ADGR* genes, mRNA expression would be disrupted in an OS cycle. This hypothesis was tested on endometrial biopsies collected in the secretory phase from prospective cohorts of women in natural and OS cycles. Consistent with the retrospective dataset, our data revealed that members of the *ADGR* gene family are expressed in the secretory phase of the natural menstrual cycle and for the first time, we show that their expression is altered by ovarian stimulation.

## INTRODUCTION

Approximately 10% of women of reproductive age in the United States suffer from infertility and a large number of these women will undergo in vitro fertilization (IVF) to conceive (Centers for Disease Control and Prevention 2018). The use of IVF has become increasingly popular, with the number of annual cycles doubling in the last decade, from 150,000 cycles per year in 2010 to more than 300,000 cycles per year in 2019 (Centers for Disease Control and Prevention 2021). Despite this increase, the live birth rate (LBR) has only increased by approximately 5%, from 32% in 2010, to 37.2% in 2019 (Centers for Disease Control and Prevention 2021).

Successful embryo implantation depends on the coordinated development of the embryo and endometrium so that when the implantation-competent embryo arrives at the endometrium, the endometrium is implantation-receptive. The period of opportunity to implant (called the window of implantation) lasts only 30-36 hours and occurs 6 to 9 days following the surge of luteinizing hormone (LH) in a natural cycle, or 4 to 7 days following progesterone administration in a hormone replacement cycle (Simon et al. 2020). Under the tight control of estradiol (E2) and progesterone (P4), the endometrium develops a secretory phenotype shortly after ovulation. Then, through the actions of P4 and cAMP in the mid-secretory phase, the endometrium decidualizes giving rise to the predecidua that regulates embryo implantation, survival and nourishment (Gellersen and Brosens 2014). Following implantation, the decidua that is associated with pregnancy regulates trophoblast invasion and the development of the placenta. Consequently, defective decidualization compromises placenta formation and/or function and is associated with fetal growth restriction and preeclampsia, which at times necessitate iatrogenic preterm birth (Garrido-Gomez et al. 2017; Turco and Moffett 2019).

Ovarian stimulation (OS), utilized for the development of multiple ovarian follicles for IVF, induces supraphysiologic levels of E2 and an early rise in P4 (Kalakota et al. 2022). These hormonal changes induce morphological, biochemical, and functional genomic modifications that disrupt receptivity and decrease implantation rates following fresh embryo transfers (Kalakota et al. 2022). Thus, with fresh embryo transfers, when the cultured embryo is returned to the uterine cavity, it may encounter a non-receptive endometrium, resulting in implantation failure. Alternatively, the embryo may encounter an endometrium that is receptive with impaired decidualization, resulting in successful embryo implantation, but suffering from the “ripple effect” of placental insufficiency and pregnancy complications (Rabaglino and Conrad 2019; Conrad 2020).

To improve pregnancy rates and reduce the risk of pregnancy complications that are more prevalent with IVF, it is crucial to advance our molecular understanding of endometrial receptivity and decidualization. Previous studies, including those from our laboratory, have shown that in the mouse, uterine G protein-coupled receptors (GPCRs) are critical regulators of endometrial receptivity and decidualization (Schaefer et al. 2021; de Oliveira et al. 2019; Radovick and Babwah 2019; Babwah 2015; Fayazi et al. 2015; Parobchak et al. 2020; Calder et al. 2014; Mohri et al. 2010; Sone et al. 2013; Kida et al. 2014; Diao et al. 2015; Verma and Arora 1990; Leon et al. 2016). In women, GPCRs might also regulate the acquisition of endometrial receptivity; this is based on their differential expression between the pre-receptive and receptive period in the secretory endometrium (Diaz-Gimeno et al. 2011). GPCRs are transmembrane proteins that regulate almost every cellular and physiologic process, including responses to gonadotropin releasing hormone (GnRH), follicle stimulating hormone (FSH) and LH (Babwah 2015). Given their ubiquitous nature, these receptors are targets of approximately 35% of all drugs approved by the United States Food and Drug Administration (Babwah 2015).

To further understand the roles of endometrial GPCRs in the regulation of decidualization we identified all GPCRs expressed across the peri-ovulatory (PO) to mid-secretory (MS) phase in natural and OS cycles. Among these GPCRs, 26 members of the adhesion G protein-coupled receptor (ADGR) family were observed to be expressed. ADGRs represent the second largest GPCR subfamily with at least 32 members distributed over nine subfamilies (Hamann et al. 2015). Recent studies suggest that ADGRs are involved in both mouse and human reproduction. In mice, ADGRD1 is needed for embryo transit from the oviduct, while ADGRG2 (GPR64) is involved in fluid reabsorption and sperm maturation in the mouse epididymis (Bianchi et al. 2021). In women, the development of the receptive endometrium is associated with the reduced expression of endometrial *ADGRG2* (Diaz-Gimeno et al. 2011; Yoo et al. 2017) while in mice it is associated with the increased expression of both ADGRG2 mRNA and protein (Yoo et al. 2017). In mice, ADGRG2 protein is also observed in the developing decidua and in human endometrial stromal cells (HESCs) and *ADGRG2* gene expression is reported to be required for HESC decidualization in vitro (Yoo et al. 2017).

Given the potential importance of endometrial ADGRs in decidualization and embryo implantation, we studied their expression across the normal menstrual cycle in a retrospective cohort of normo-ovulatory women (Talbi et al. 2006) and found that many *ADGRs* are differentially expressed between the proliferative and secretory phase of the cycle. This strongly suggests that in the human endometrium, *ADGRs* are E2- and/or P4-regulated genes. This is supported by mouse data that showed *Adgrg2* mRNA expression is P4-dependent (Yoo et al. 2017). Together, these findings lead us to hypothesize that in women, *ADGR* gene expression would be disrupted in an OS cycle. Based on their expression profiles, we further hypothesize that in addition to *ADGRG2*, other *ADGR* genes regulate HESC decidualization. These hypotheses were tested on endometrial biopsies from prospective cohorts of women in natural and OS cycles and the uterus from pregnant and non-pregnant mice; the findings from our studies are reported herein.

## MATERIALS & METHODS

### *In silico* Analysis

An *in silico* analysis was performed using GEO dataset GDS2052, which consists of the microarray analysis of human endometrial gene expression across the menstrual cycle (Talbi et al. 2006). GDS2052 includes 27 normo-ovulatory women ranging from 23 to 50 years old, with regular menstrual cycles lasting 24 to 35 days (Figure 1). Women with laparoscopy proven endometriosis or inflammation within the endometrium at the time of the biopsy, were excluded from the study. Endometrial tissue was collected from either hysterectomy samples, performed for benign indications (n=20), or endometrial biopsies (EMBs) obtained via aspiration with a Pipelle catheter or endometrial curetting (n=7). Pathological diagnoses at the time of hysterectomy uncovered uterine fibroids (n=13), adenomyosis (n=3), ovarian cyst (n=1) and pelvic organ prolapse (n=3). Subjects recruited for EMB were volunteers who had not used steroid medication for at least three months prior to the biopsy. Endometrial tissue was collected at different phases of the menstrual cycle and staged in a blinded fashion by two or more pathologists using Noyes criteria (Noyes, Hertig, and Rock 1975) for menstrual cycle phase determinations.

**Figure 1.**
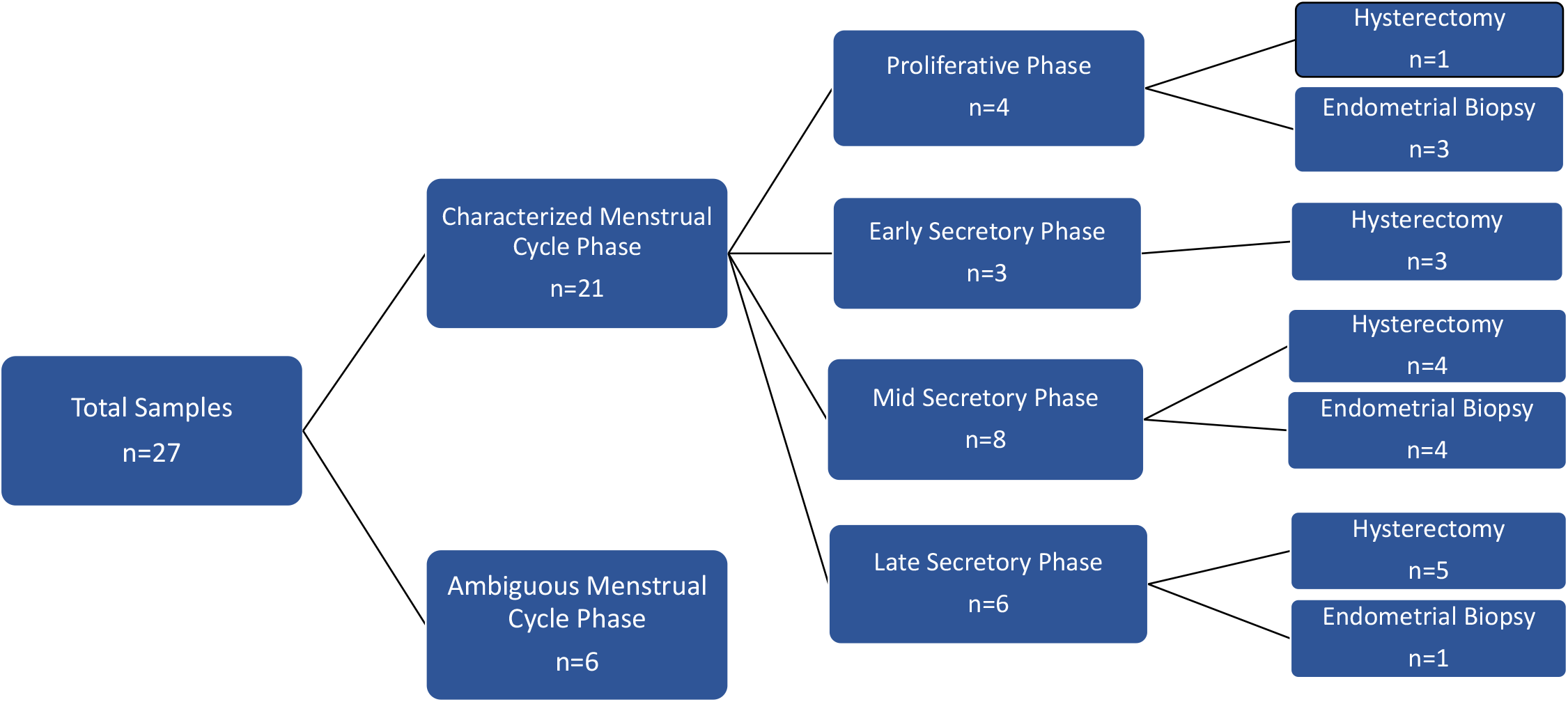
Flow chart illustrating human study groups selected from Dataset Record GDS2052 (Endometrium throughout the menstrual cycle) for in silico analysis.

For the in silico analysis reported in this study, 21 subjects were included, while six subjects with an ambiguous cycle stage were excluded (Figure 1; Supplemental Table 1). The subjects included, ranged from 23-49 years old. Staged endometrial samples included proliferative (n=4), early secretory (n=3), mid secretory (n=8) and late secretory (n=6) phases. Endometrial *ADGR* gene expression in these 21 samples was determined using the high-density human genome (HG) U133 Plus 2.0 Arrays (Affymetrix), containing 42,203 genes and 12,397 expressed sequence tags (ESTs). This array contains probes against all 32 *ADGR* genes known to encode protein products (Hamann et al. 2015) (Supplemental Table 2). Probe level intensities were processed and hit, or raw counts were normalized as described (Talbi et al. 2006). Differentially expressed genes were identified using limma/voom (v3.52.2) and the number of significant, differentially expressed genes was determined using 5% false detection rate (FDR) (q-value <0.05) as the threshold.

### Prospective Tissue and Serum Collection

#### Endometrial Biopsy Sample Collection

EMBs were prospectively collected from women in natural and ovarian stimulation cycles using an Institution Review Board approved protocol (Pro2018002041) at Rutgers New Jersey Medical School and a university affiliated fertility clinic. For those women who underwent sampling in a natural menstrual cycle, inclusion criteria included (1) age between 21-45; (2) regular menstrual cycle (21-35 days); (3) prior pregnancy. Exclusion criteria included: (1) endocrine or autoimmune disorders; (2) anatomic disorders of the reproductive tract, including hydrosalpinges, submucosal fibroids and endometrial polyps; (3) recurrent pregnancy loss; (4) use of hormonal medication for at least 3 months prior to enrollment; (5) history of pregnancy complications such as preeclampsia or intrauterine growth restriction; (6) pregnant or trying to conceive. A total of 15 women were recruited in the natural cycle from ages 21 to 33 years old (Supplemental Table 3). EMBs were collected in the proliferative (PRO) phase (cycle days 10-13; n=4) and in the peri-ovulatory (PO) period and mid-secretory (MS) phase in natural cycles using a disposable endometrial Pipelle (MedGyn Pipette). Women recruited for secretory phase biopsies, used Pregmate ovulation test strips ovulation predictor to detect the urinary LH surge prior to ovulation. The day of LH surge detection was deemed LH+0, and biopsies were either performed in the PO period (PO; n=6) (henceforth referred to as PO-NC) or the mid-secretory (LH+9; n=5) (henceforth referred to as MS-NC) phase (Figure 2). Women recruited in the PO and MS periods were asked to present themselves for a biopsy two or nine days after detecting a rise in urinary LH. Since the LH surge lasts approximately 12-24 hrs, depending on when in that timespan the surge was detected, the PO and MS phase biopsies were anywhere in a window corresponding to LH+1-2 and LH+8-9, respectively. Since ovulation occurs about 36 hrs after the onset of the surge, in the LH+1-2 period, sampling could have been before or after ovulation. Hence, we refer to this as the PO period. The median ages for the PRO phase, PO period and MS phase were 27.5, 29.5, and 28 years old respectively (Supplemental Table 3, Supplemental Table 5).

**Figure 2.**
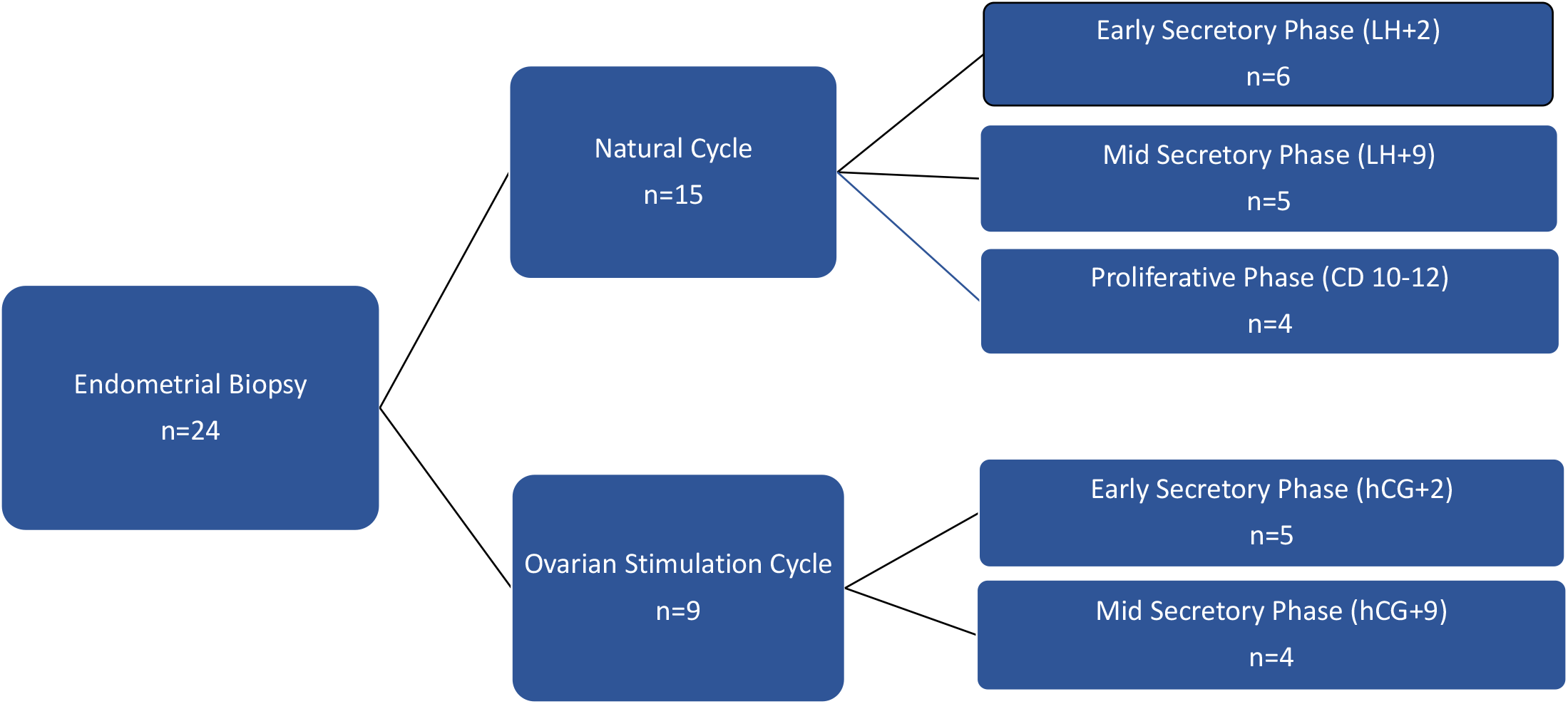
Flow chart illustrating human study groups from whom endometrial biopsies were collected for RNA sequencing analysis.

Women undergoing OS were recruited from a university affiliated fertility clinic. Inclusion criteria were (1) age between 21-45; (2) IVF without fresh embryo transfer; (3) diagnosis of male factor, tubal factor, or unexplained infertility; (4) undergoing OS for oocyte cryopreservation; (5) plan to freeze all oocytes or embryos obtained during the treatment. The same exclusion criteria applied to women in natural cycles were applied to women undergoing OS. OS was performed using a GnRH-antagonist (Cetrotide, Ganirelix or Fyremadel) protocol without embryo transfer. Gonadotropins used for stimulation include Menopur used with a combination of Follistim, Gonal-F or low dose HCG. During stimulation, patients were monitored regularly using ultrasound and serum hormone levels. Once follicles reached 17-19 mm, final oocyte maturation was triggered with hCG (10,000 units). Nine women were recruited, ages ranging from 31-42 years old (Supplemental Table 4) The EMB samples were obtained following the hCG trigger, in either the early secretory phase (hCG+2; n=5) (henceforth referred to as PO-OS) or the mid-secretory phase (hCG+9; n=4) (henceforth referred to as MS-OS), using a disposable endometrial Pipelle (MedGyn Pipette) (Figure 2). Since women are biopsied relative to the time of the hCG injection, biopsies at hCG+2 and hCG+9 are more precisely timed compared to the PO and MS phase biopsies in natural cycles. Median ages for hCG+2 and hCG+9 were 33 and 37.5 years old respectively (Supplemental Table 4, Supplemental Table 5).

#### Histological Analysis

Following EMB collection, tissue samples were fixed in 10% neutral buffered formalin solution, paraffin-embedded and sectioned at 7 μm. Sections were then deparaffinized, rehydrated, and stained with hematoxylin and eosin (H&E). Two pathologists, blinded to the treatment groups, independently determined endometrial dating on H&E stained sections. Histologic summary confirmed menstrual cycle stage for all samples included in the analysis (data not shown).

#### Analysis of Estradiol and Progesterone

At the time of EMB, blood was collected for serum measurements of E2 and P4 levels. An electrochemiluminescence immunoassay (ECLIA) was utilized to measure both serum hormones (E2: Labcorp test: 004515 and P4: Labcorp test: 004317). To compare serum hormone levels non parametric statistical analyses were performed after normality was assessed using a Schapiro Wilk’s Test. Values were expressed as medians with interquartile ranges to assess the spread of the data. (Supplemental Table 5).

#### RNA Extraction and Next Generation Sequencing

Total RNA was extracted from the EMBs using RNeasy Plus Mini Kit (Qiagen, Frederick, MD, USA) according to the manufacturer’s protocol. Following extraction, the RNA concentration and integrity were determined. RNA samples were then sent to the Computational Genomics Core at Albert Einstein College of Medicine, Bronx, NYC, for next-generation mRNA sequencing. The raw transcriptome reads were assessed for quality control (FASTQC v 0.11.8) and trimmed for quality/adapter contamination (cutadapt v2.5). Trimmed reads were aligned to the human genome (GRCh37) using STAR (v2.6.1), followed by transcript abundance calculations and hit count extraction with StringTie (v2.0) and featureCounts (v1.6.4) respectively. Hit count normalization and differential gene expression group cross comparisons were performed using DESeq2 (v1.26.0). Significant differentially expressed gene thresholds were set to FDR adjusted p<0.05.

#### qPCR Analysis of Human *ADGR* Gene Expression

Total RNA (0.5 μg) was extracted using the RNeasy Mini Kit (Qiagen, Germantown, MD, USA) from EMBs and reverse transcribed using qScript cDNA Supermix (Quantabio, Beverly, MA, USA). Gene expression was evaluated using human gene-specific primers (Supplemental Table 6). qPCR was conducted using the QuantiNova SYBR Green PCR kit (Qiagen, Frederick, MD, USA).

#### Mouse Studies

Animal studies were approved by the Institutional Animal Care and Use Committee at Rutgers New Jersey Medical School. All data presented in this study were generated using C57BL/6J wild type mice (Jackson Laboratories). Studies involving non-pregnant mice: Random cycling females were euthanized at 10 weeks of age. The uterus was removed and preserved in RNA later (Qiagen). Studies involving pregnant mice: Females (8-10 weeks old) were mated with stud males, and the day of mating was designated as day 0, with embryonic day (E) 0.5 being noon on the day a mating plug was observed. Mice were then euthanized on E0.5, E5.5, E6.5 and E7.5. On E0.5, the entire uterus was isolated and used for subsequent analysis, while on E5.5, E6.5 and E7.5, the myometrium was removed to isolate the underlying decidua. Whole uteri from E0.5, as well as decidua from E5.5, E6.5 and E7.5 were preserved in RNA later.

#### RT-PCR and q-PCR Analysis of Mouse *Adgr* Gene Expression

RT-PCR was performed on whole uteri from non-pregnant dams and decidua from pregnant dams at E7.5. qPCR was performed on whole uteri from pregnant dams at E0.5 and decidua on E5.5 and 6.5. For both RT-PCR and qPCR, total RNA (0.5 μg) was extracted and reverse transcribed using qScript cDNA Supermix (Quantabio, Beverly, MA, USA). Gene expression was evaluated using mouse gene-specific primers (Supplemental Table 7). For RT-PCR, PCR products were visualized on an agarose gel after 30 cycles of amplification. qPCR was conducted using the QuantiNova SYBR Green PCR kit (Qiagen, Frederick, MD, USA).

#### qPCR Analysis (Human and Mouse)

All measurements were performed in triplicate and the gene expression level was determined as a mean of the triplicate. Relative gene expression was determined using the 2^-ΔΔCt^ method (Livak and Schmittgen 2001) and expressed as fold change normalized to the human RNA18SN5 or mouse *Rn18s* housekeeping genes. Fold changes were then compared using a Kruskal-Wallis test for three or more samples with data expressed in mean rank. For two samples, group comparisons were made using the Mann-Whitney U test and data were expressed as medians with interquartile range (IQR). Statistical analyses were performed using Prism Version 9.0 (GraphPad). Statistical significance was defined as p<0.05.

#### Tissue Culture of Primary Human Endometrial Stromal Cells (HESC)

Three primary cell lines were established from two proliferative phase endometrial samples according to Chen et al (Chen and Roan 2015). The cell lines were cultured in phenol red-free DMEM F12 (Gibco Cat# 11039-021) supplemented with 10 ng/ml BFGF (EMD Millipore Cat# GF003), 1 mM sodium pyruvate (Sigma Cat # D2906) penicillin-streptomycin (ThermoFisher Cat # 15070063) and 10% charcoal-stripped fetal bovine serum (ThermoFisher Scientific Cat # A3382101). This medium is referred to as basal growth medium.

#### *In Vitro* Decidualization of Primary HESC

Primary HESCs cultured in a basal growth medium and induced to undergo decidualization by supplementing the medium with 10 nM E2, 1 μM P4 and 50 μM db-cAMP (Michalski et al. 2018). Cells were subjected to *in vitro* decidualization for three and six days. Decidualization was confirmed visually by cellular morphological changes from a fibroblastic to epithelioid phenotype, and quantitatively via increased expression of *IGFBP1, PRL* and *FOXO1* mRNA

## RESULTS

### In silico analysis of *ADGR* Gene Expression in a Natural Menstrual Cycle

To quantify endometrial *ADGR* gene expression across the natural menstrual cycle, an in silico analysis was performed using an existing GEO dataset (GDS2052). GDS2052 is derived from the microarray analysis of human endometrial gene expression throughout the menstrual cycle. The heatmap shows that all 32 known *ADGR* coding genes, representing 9 ADGR subfamilies were expressed across the menstrual cycle (Figure 3A). Of the genes identified, 13 ADGRs were noted to have multiple probes per gene (Supplemental Table 2), and overall trends of directionality in gene expression changes were conserved across the various probes (data not shown). *ADGR* gene expression (based on normalized hit counts) across the proliferative (PRO) and early, mid and late secretory phase (ES, MS and LS, respectively) of the menstrual cycle was visualized on a heatmap (Figure 3A) and differential gene expression was performed by conducting pairwise comparisons of normalized hit counts (Figure 3B-Q). When comparing the PRO versus ES, MS and LS phases, 1, 3 and 11 differentially expressed ADGRs were identified, respectively and these represented a total of 12 *ADGR* genes (Figure 3; Supplemental Table 8). Next, a comparison of gene expression was made within the secretory phase. In the secretory phase, 3 ADGRs were found to be differentially expressed between the ES and MS phases (Figure 3; Supplemental Table 9), while 11 were differentially expressed between the ES and LS phases (Figure 3; Supplemental Table 9). Together, these data show that *ADGR* gene expression varies significantly with menstrual cycle stage, suggesting that *ADGR* expression is regulated by E2 and P4. The data also suggest that *ADGRB2, ADGRF1*, and *ADGRL1*, which were upregulated in the MS phase (corresponding to the window of implantation) relative to the PRO phase (Figure 3; Supplemental Table 8) regulate predecidualization.

**Figure 3.**
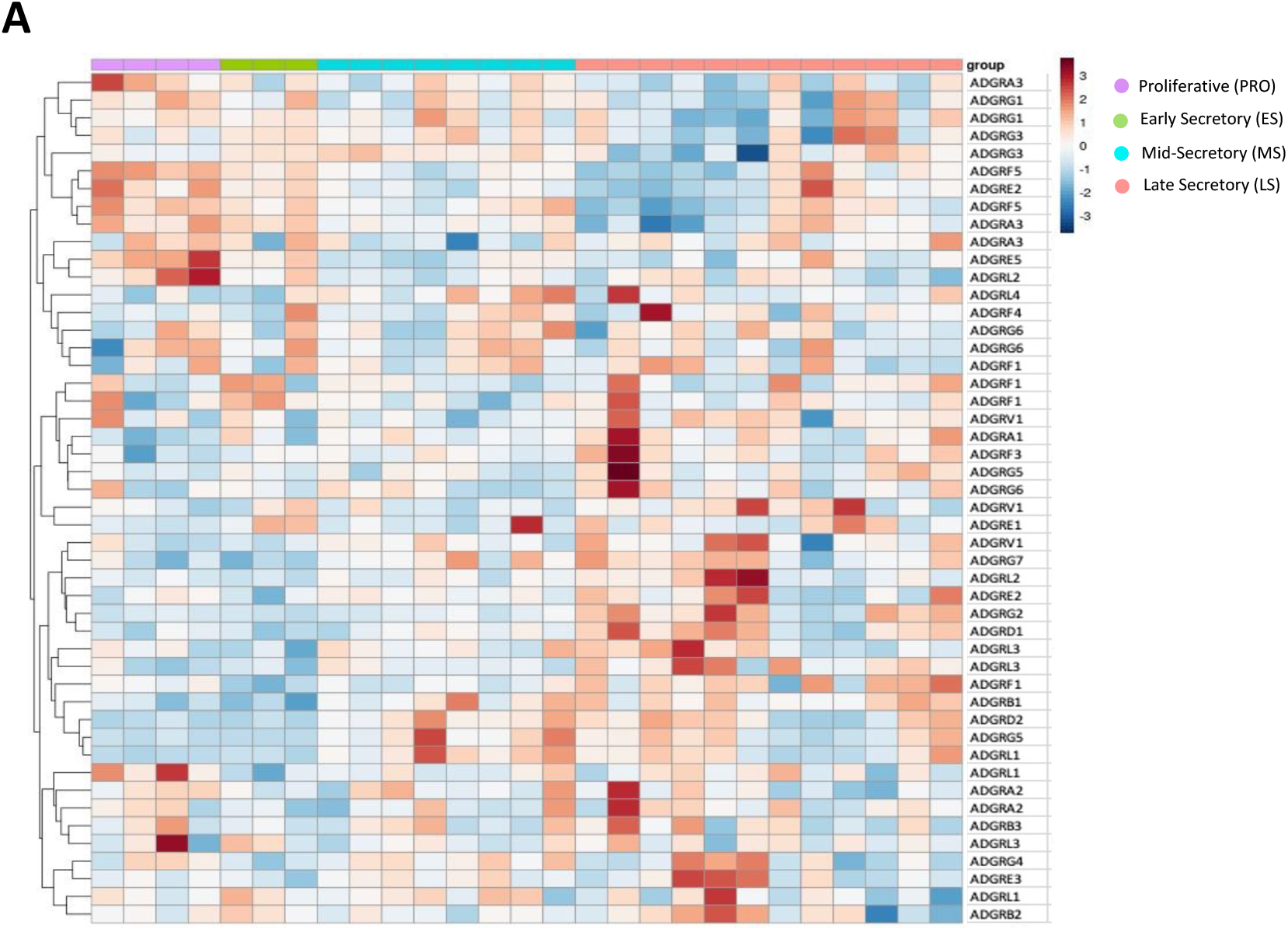
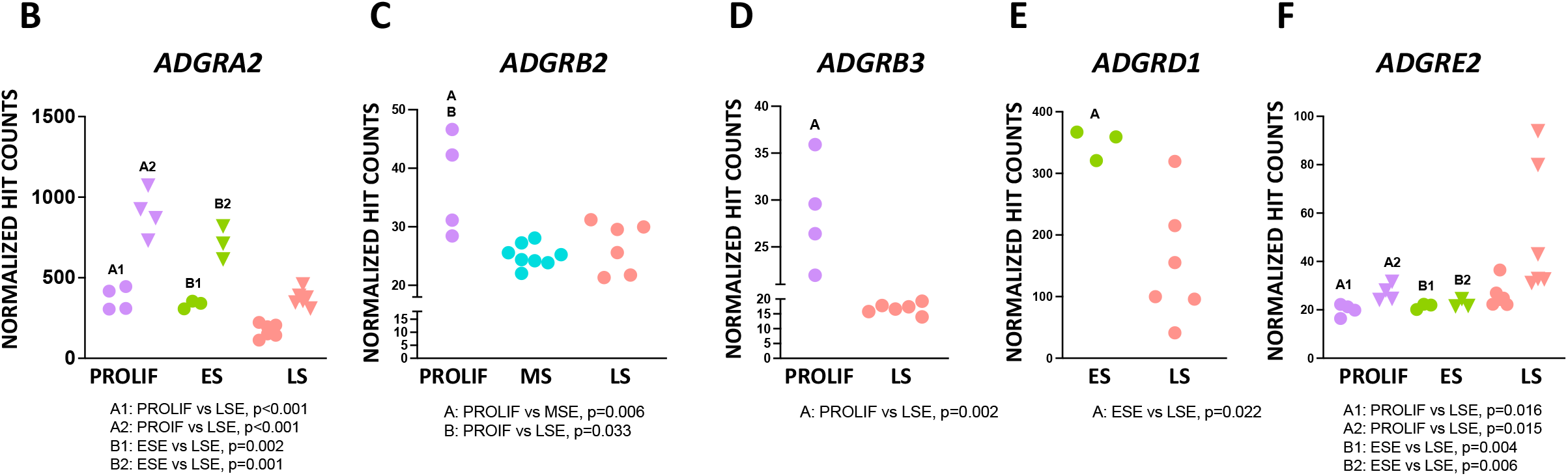
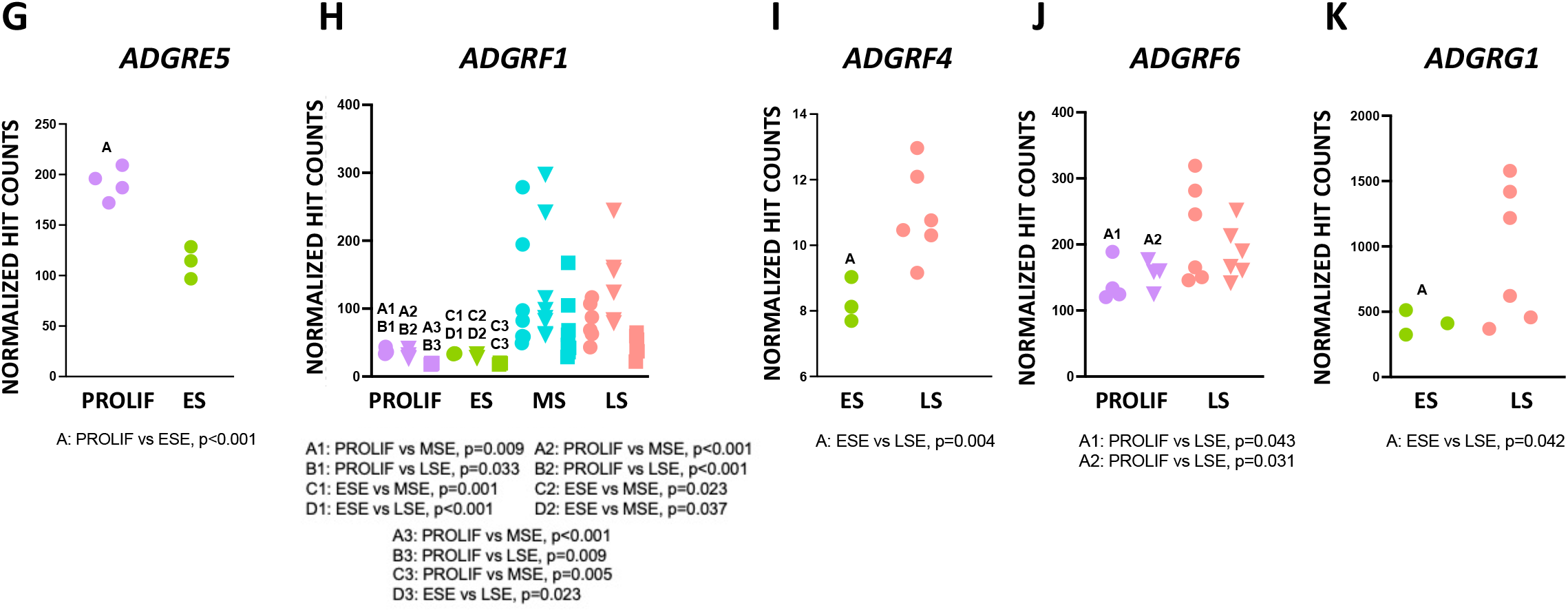
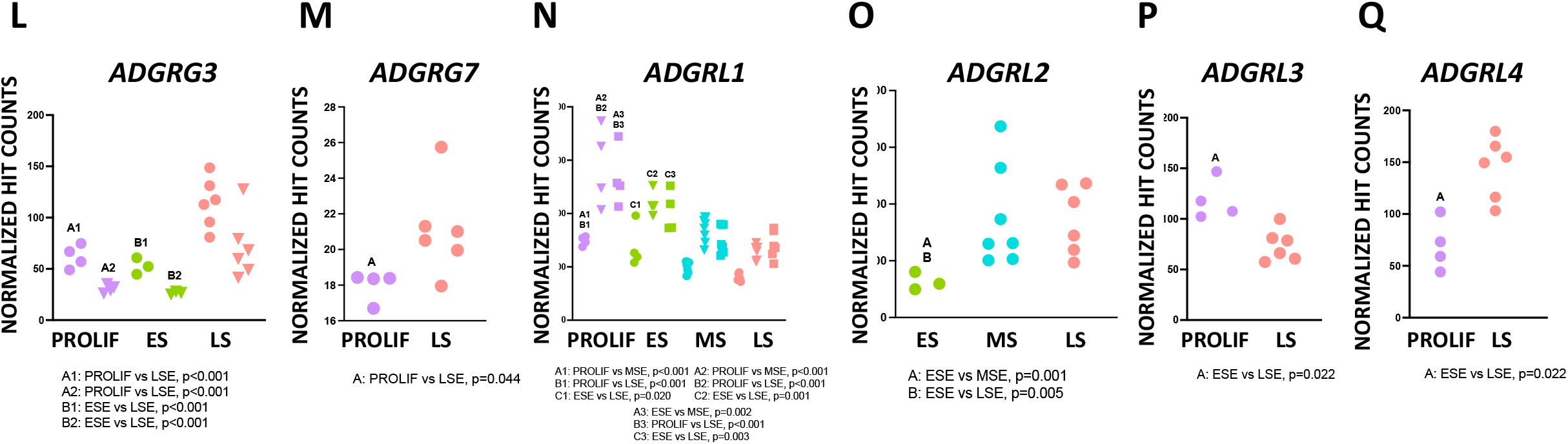
In silico analysis of ADGR mRNA expression across the menstrual cycle. (A) Heat map of *ADGR* expression among subjects from the proliferative and secretory phase. (B) Differentially expressed *ADGR* genes between the proliferative vs early and mid-secretory phase. Gene expression levels calculated as mean normalized hit counts. Differential gene expression group cross-comparisons were performed using DESeq2 (v1.26.0). Significant differentially expressed gene thresholds were set at FDR adjusted p <0.05.

### Baseline Characteristics of Prospective Natural Cycle (NC) and Ovarian Stimulation (OS) Cohorts

To understand the impact of OS on *ADGR* expression and the functional consequences on the development of the endometrium during the secretory phase, biopsies were prospectively collected from women in natural and OS cycles. Following screening, nine healthy eumenorrheic women, six with PCOS, and six with HA were recruited to the study. Baseline characteristics and baseline reproductive hormone profiles of the study participants are presented in Supplemental Table 1. An analysis of the data revealed there was no significant difference in age, weight, body mass index (BMI), baseline FSH, E2 or SHBG levels between the three groups of women. However, serum LH and serum anti-Müllerian hormone (AMH) levels were significantly higher in women with PCOS (P=0.003 for LH, and P=0.005 for AMH).

### Prospective analysis of *ADGRB2, ADGRF1*, and *ADGRL1* mRNA Expression in the Proliferative and Secretory Phase in a Natural Menstrual Cycle

Based on the in silico analysis of GEO dataset GDS2052, given the potential role of *ADGRB2, ADGRF1*, and *ADGRL1* in endometrial decidualization, we validated their expression in prospectively collected endometrial biopsies in the PRO (n=4) and MS (n=5) phase (Figure 2) of the natural menstrual cycle. Since *ADGRG2* was reported to enhance stromal cell decidualization (Yoo et al. 2017), it was also included in this analysis. Based on qPCR analysis, it was observed that while *ADGRG2* increased significantly in expression in the MS phase, *ADGRB2, ADGRF1* and *ADGRL1* showed no significant change in expression (Figure 4).

**Figure 4.**
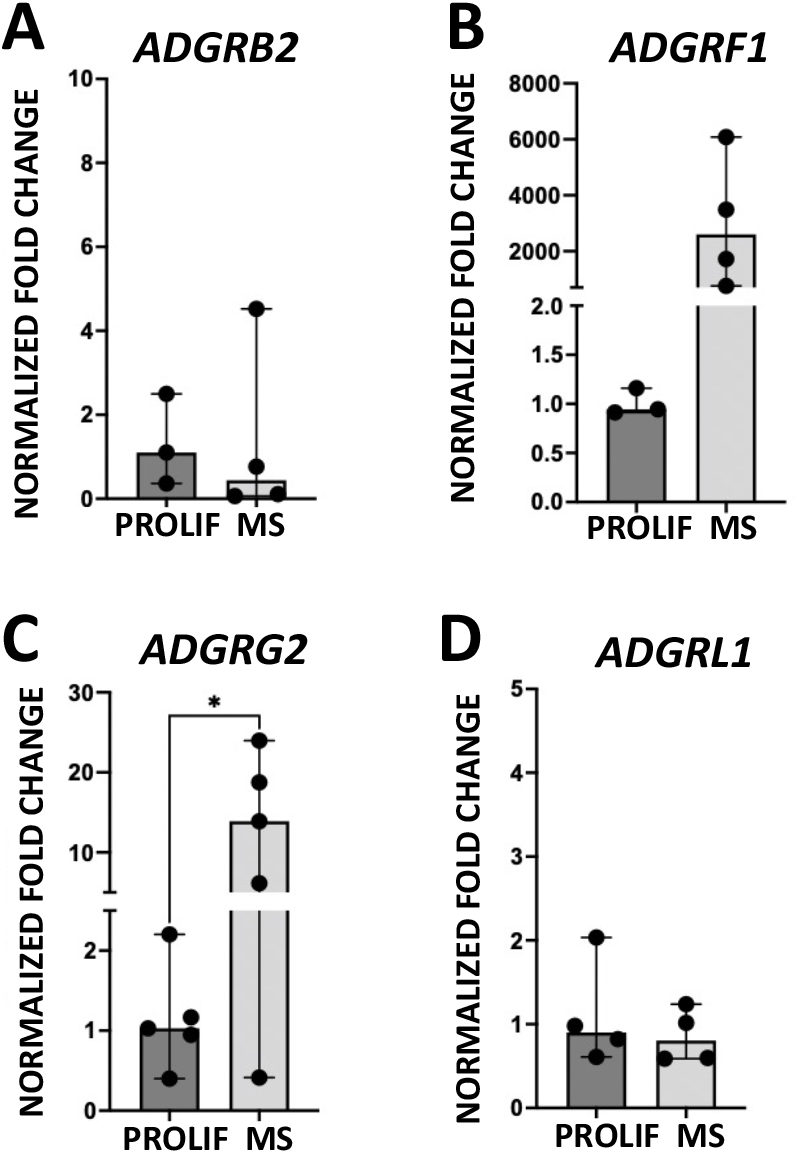
*ADGRB2, ADGRF1, ADGRG2* and *ADGRL1* mRNA expression in the human endometrium in the proliferative and MS phase. MS: mid-secretory.

### Prospective analysis of *ADGR* mRNA Expression in the Secretory Phase in Natural Menstrual and Ovarian Stimulation Cycles

Prospectively collected endometrial tissue samples from the PO-MS period of the natural and ovarian stimulation cycles were analyzed using mRNA sequencing (mRNA seq) to determine differential gene expression. To determine whether *ADGR* genes were among the DEGS, mRNA seq data was examined. This first revealed that among the natural and ovarian stimulation cycles spanning the PO to MS phase, 26 coding *ADGRs*, representing 8 of the 9 known subfamilies were expressed (Figure 5A). Expression of *ADGRA1, ADGRC1, ADGRC2, ADGRC3, ADGRD2* and *ADGRG4* was not detected.

**Figure 5.**
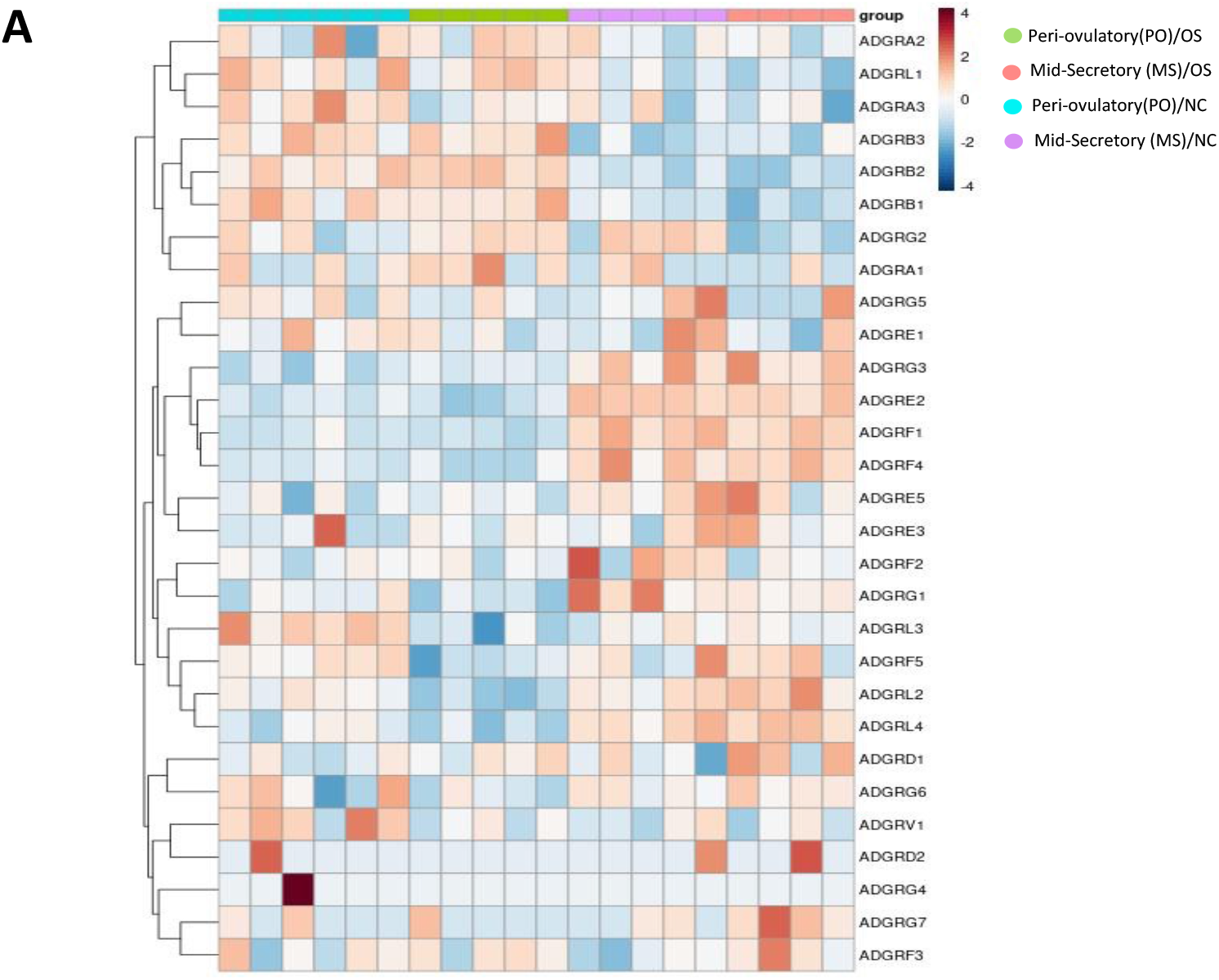
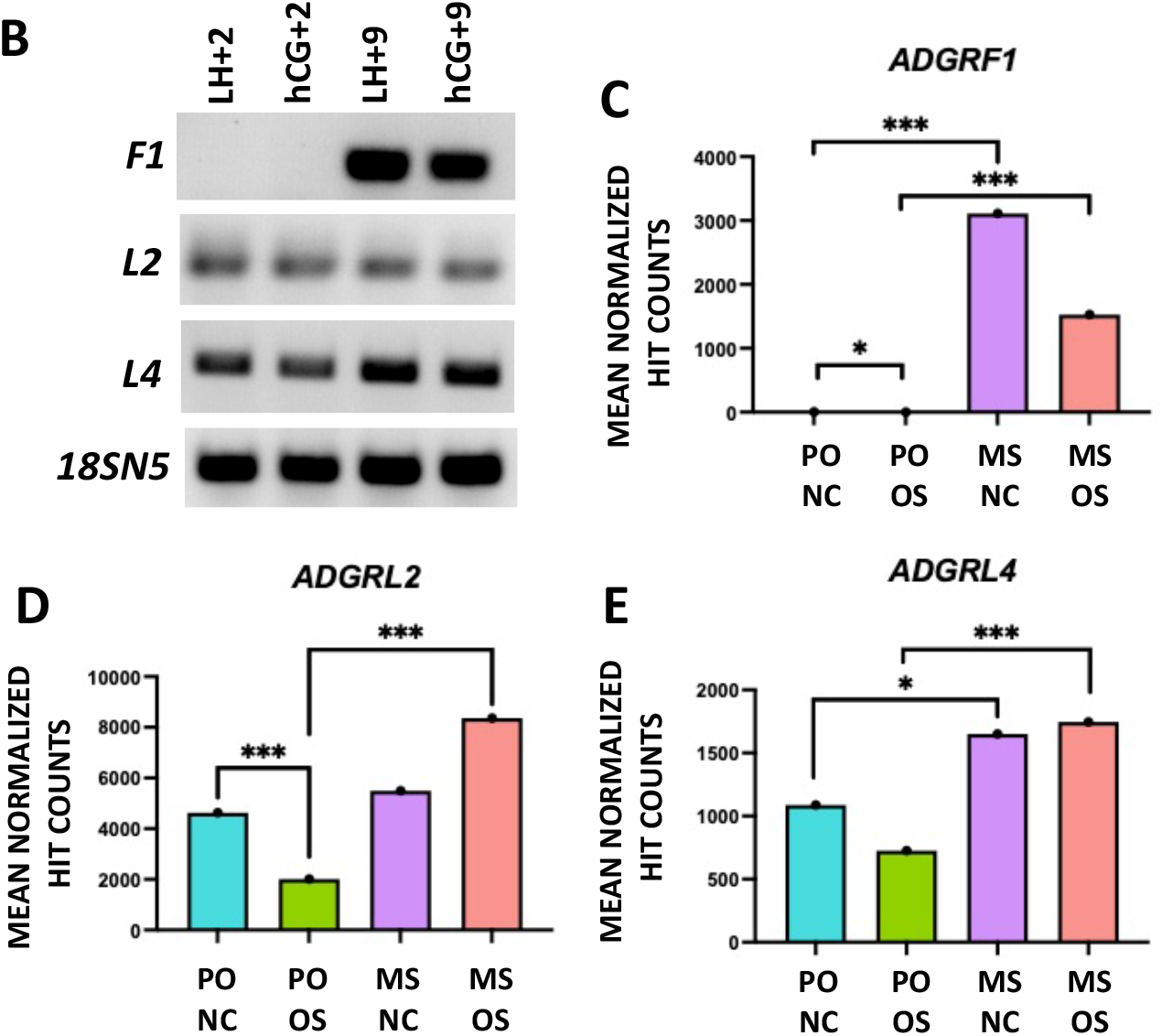
RNA seq analysis of *ADGR* mRNA expression in the PO and MS phase of natural and ovarian-stimulated cycles. (A) Heat map of *ADGR* expression among patients from the PO and MS phase of natural and OS cycles. (B-E) *ADGRF1, ADGRL2* and *ADGRL4* mRNA expression in the PO and MS phase of natural and OS cycles. Significant differentially expressed gene thresholds were set at FDR adjusted p <0.05. NC: natural cycle, OS: ovarian-stimulated cycle, MS: mid-secretory, PO: peri-ovulatory.

The three DEGs *(ADGRF1, ADGRL2*, and *ADGRL4)* identified in the in silico analysis when comparing the ES to MS phase (Figure 3; Supplemental Table 8) were also expressed by RT-PCR in the prospectively collected biopsies spanning the PO-MS period of natural and OS cycles (Figure 5B). These genes were further analyzed using the RNA seq dataset. When comparing the MS phase to the PO period of a NC, based on normalized expression counts, of these three genes, both *ADGRF1* and *ADGRL4* were found to be upregulated while *ADGRL2* was unchanged in expression (Figure 5C; Supplemental Table 10). However, when comparing the MS phase to the PO period in an OS cycle, all three genes, *ADGRF1, ADGRL2* and *ADGRL4* were increased in expression (Figure 5C-E; Supplemental Table 10). When comparing the PO period in a natural vs OS cycle, both *ADGRF1* and *ADGRL2* were decreased expression, while *ADGRL4* was unchanged (Figure 5E; Supplemental Table 10). When comparing the MS phase in a natural vs OS cycle, neither *ADGRF1, ADGRL2* nor *ADGRL4* exhibited a change in expression (Figure 5C-E; Supplemental Table 10).

#### Characterization of ADGRs with Respect to Decidualization in Early Mouse Pregnancy

Given the potential roles of *ADGRB2, ADGRF1, ADGRL1, ADGRL2*, and *ADGRL4* in human endometrial decidualization and the ethical restrictions in studying the early pregnancy decidua after embryo implantation in women, a mouse model of early human pregnancy was studied to understand the role of *ADGR* genes in decidualization. To begin, an RT-PCR screen was conducted to identify which *Adgr* genes, representing subfamilies, *Adgra, Adgrb, Adgre, Adgrf, Adgrg*, and *Adgrl*, are expressed in the non-pregnant (NP) uterus and E7.5 decidua. Pregnancy day E7.5 was chosen for this study since on this day, decidua formation is complete. The five subfamilies selected for analysis were based on the differentially expressed *ADGR* genes identified in the *in silico* analysis (Figure 3). Based on the RT-PCR screen, it was observed that the following *Adgr* genes were expressed either in the NP and/or E7.5 samples: *Adgra1, Adgra2, Adgra3* (Figure 6A), *Adgre1, Adgre5* (Figure 7A),*Adgrf1, Adgrf2, Adgrf4, Adgrf5* (Figure 8A) *Adgrg1, Adgrg2, Adgrg3, Adgrg5, Adgrg6, Adgrg7* (Figure 9A), *Adgrl1, Adgrl2, Adgrl1* and *Adgrl4* (Figure 10A). Following the RT-PCR screen, those genes that were expressed in the E7.5 decidua were chosen for qPCR characterization of expression in early pregnancy at E0.5, 5.5 and 6.5. These genes are *Adgra1, Adgra2, Adgra3* (Figure 6), *Adgre1, Adgre5* (Figure 7), *Adgrf2, Adgrf4, Adgrf5* (Figure 8) *Adgrg1, Adgrg2, Adgrg3, Adgrg6, Adgrg7* (Figure 9) *Adgrl1, Adgrl2* and *Adgrl4* (Figure 10). Among these genes, *Adgra2* (Figure 6C), *Adgre1* (Figure 7B), *Adgrg1, Adgrg6, Adgrg7* (Figure 9B, E and F), *Adgrl1, Adgrl2* and *Adgrl4* (Figure 10B, C and D) all demonstrated significantly different expression between E0.5 and E5.5 or 6.5. Whereas *ADGRG2, ADGRG3* (Figure 9C and D) and *ADGRL2* (Figure 10C) only exhibited significant changes in expression between E5.5 and E6.5. *ADGRA3* (Figure 6D) and *ADGRE5* (Figure 7C) showed no significant change in expression across experimental groups.

**Figure 6.**
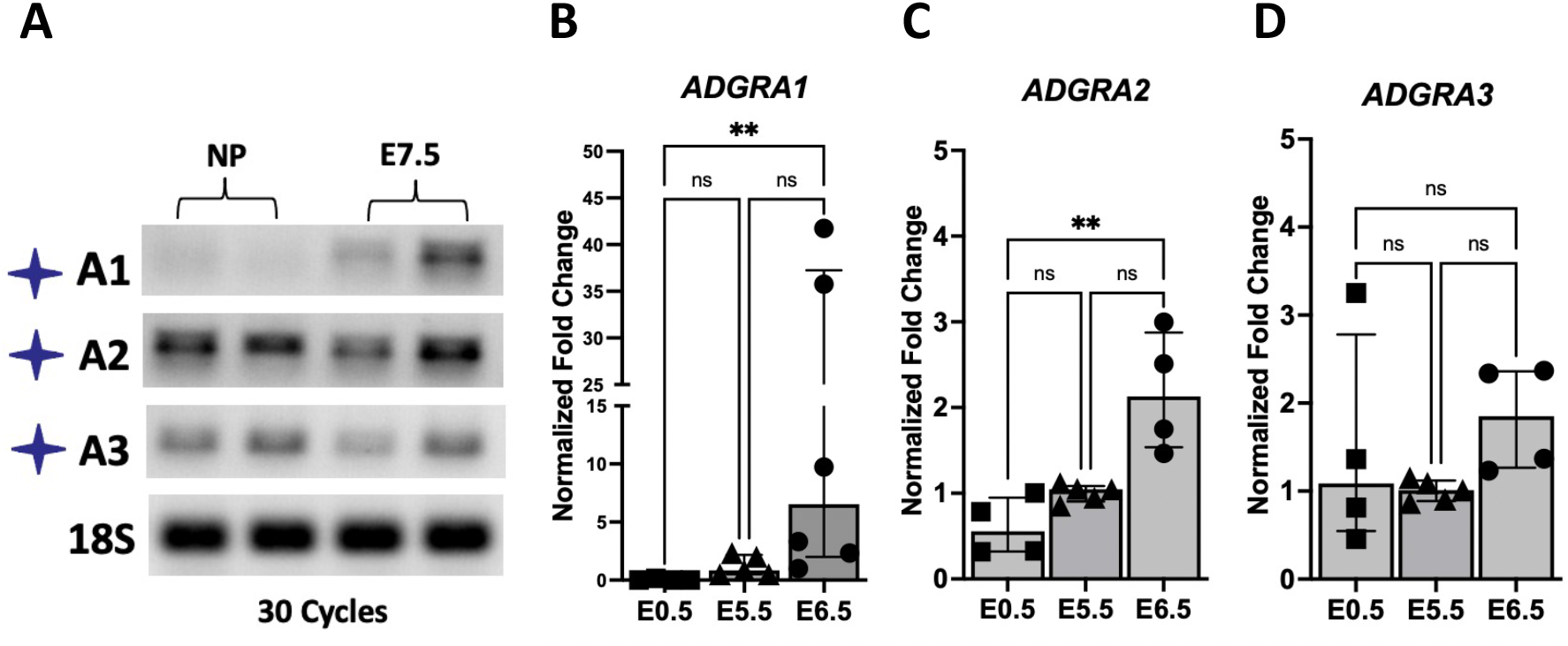
Expression of genes in the ADGRA subfamily in the non-pregnant and pregnant mouse on E7.5. (A) RT-PCR analysis of gene expression in the ADGRA subfamily. (B-D) For genes that showed expression at E7.5 based on RT-PCR analysis (A), their expression was validated and quantified further by qPCR across early mouse pregnancy at E0.5 (in the whole uterus), E5.5 (in the decidua) and E6.5 (in the decidua).

**Figure 7.**
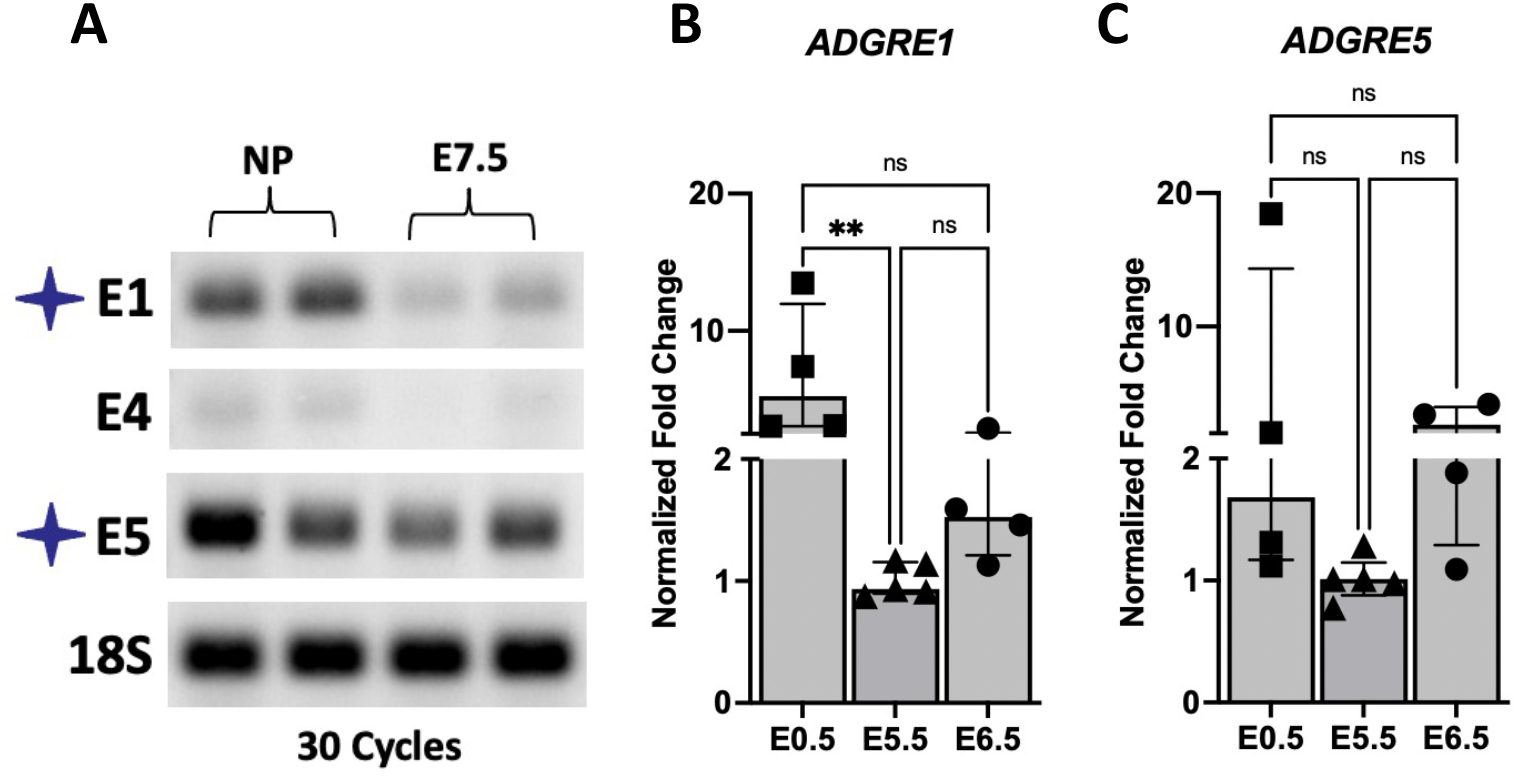
Expression of genes in the ADGRE subfamily in the non-pregnant and pregnant mouse on E7.5. (A) RT-PCR analysis of gene expression in the ADGRE subfamily. (B-C) For genes that showed expression at E7.5 based on RT-PCR analysis (A), their expression was validated and quantified further by qPCR across early mouse pregnancy at E0.5 (in the whole uterus), E5.5 (in the decidua) and E6.5 (in the decidua).

**Figure 8.**
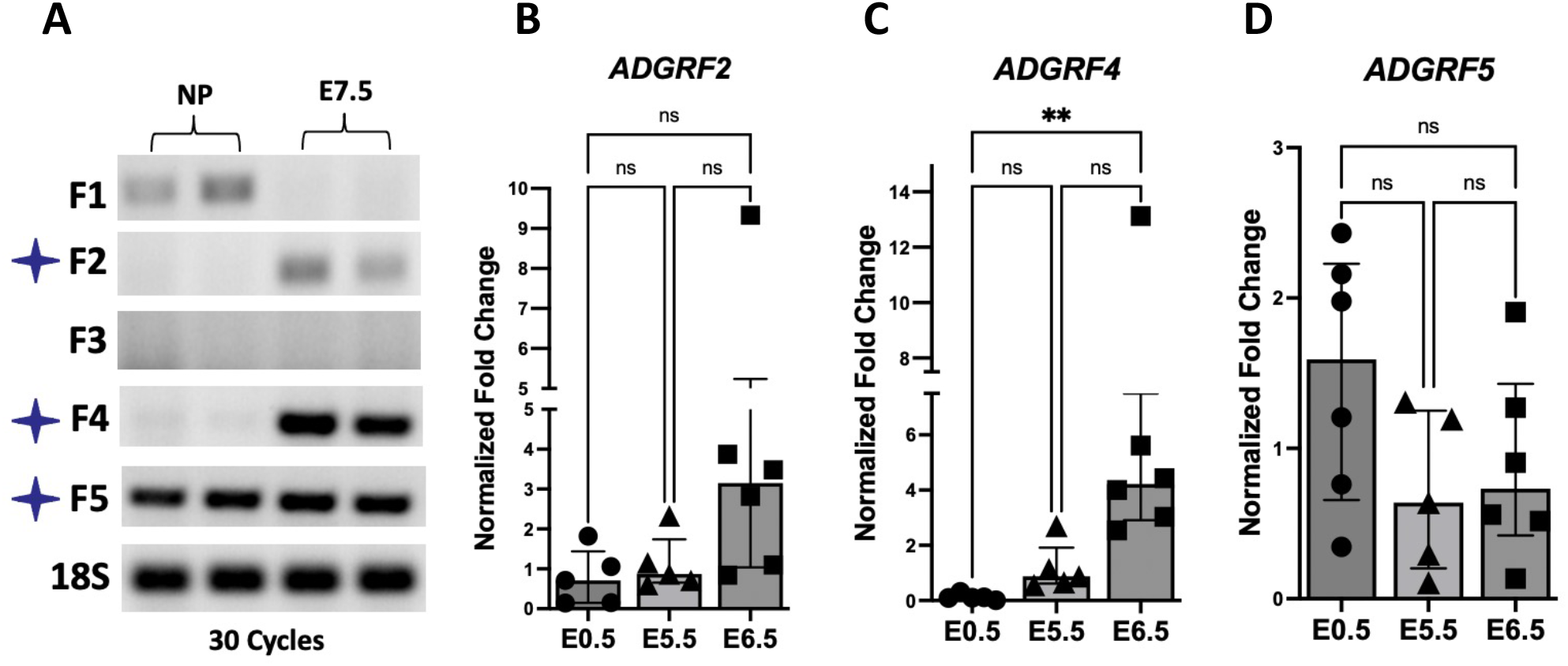
Expression of genes in the ADGRF subfamily in the non-pregnant and pregnant mouse on E7.5. (A) RT-PCR analysis of gene expression in the ADGRF subfamily. (B-D) For genes that showed expression at E7.5 based on RT-PCR analysis (A), their expression was validated and quantified further by qPCR across early mouse pregnancy at E0.5 (in the whole uterus), E5.5 (in the decidua) and E6.5 (in the decidua).

**Figure 9.**
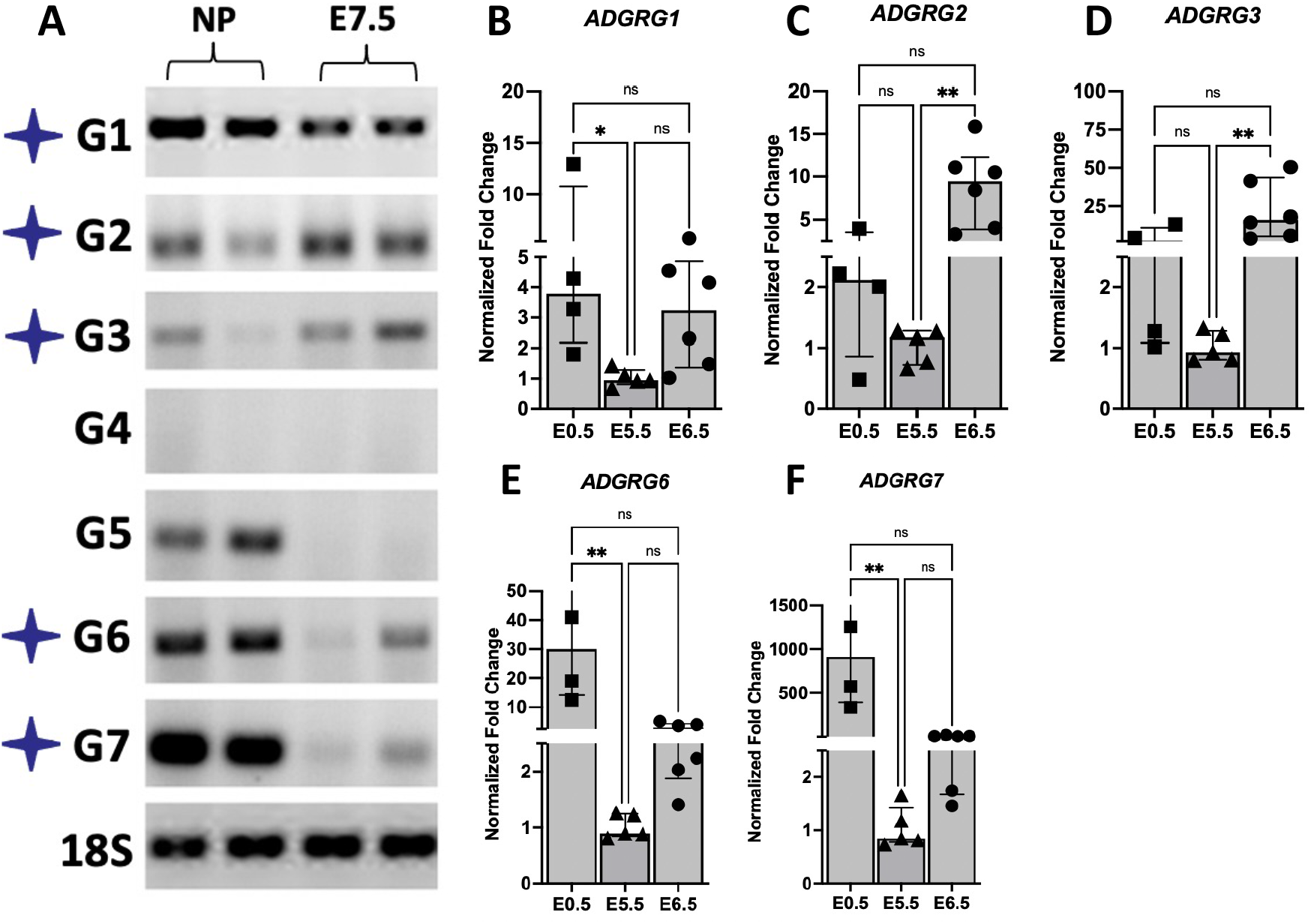
Expression of genes in the ADGRG subfamily in the non-pregnant and pregnant mouse on E7.5. (A) RT-PCR analysis of gene expression in the ADGRG subfamily. (B-F) For genes that showed expression at E7.5 based on RT-PCR analysis (A), their expression was validated and quantified further by qPCR across early mouse pregnancy at E0.5 (in the whole uterus), E5.5 (in the decidua) and E6.5 (in the decidua).

**Figure 10.**
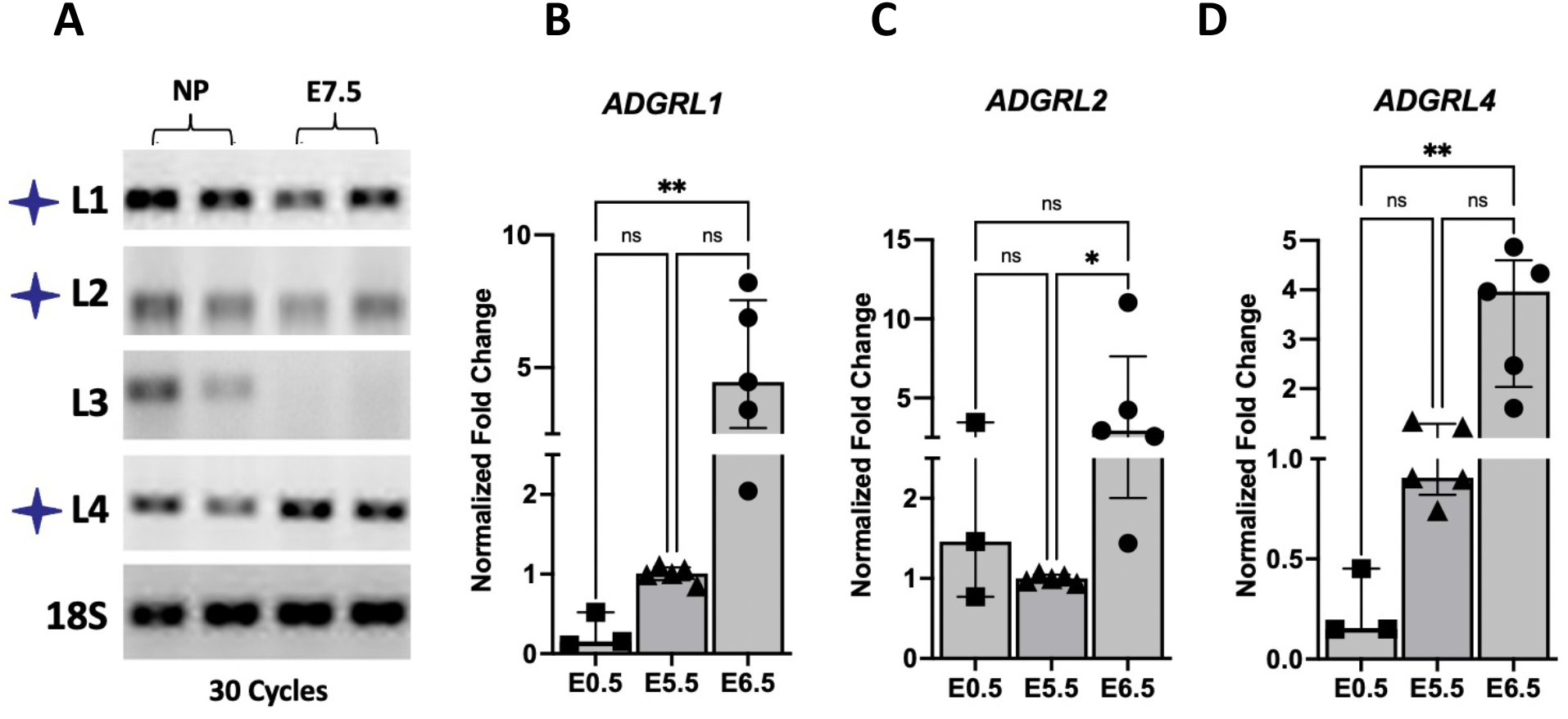
Expression of genes in the ADGRL subfamily in the non-pregnant and pregnant mouse on E7.5. (A) RT-PCR analysis of gene expression in the ADGRL subfamily. (B-D) For genes that showed expression at E7.5 based on RT-PCR analysis (A), their expression was validated and quantified further by qPCR across early mouse pregnancy at E0.5 (in the whole uterus), E5.5 (in the decidua) and E6.5 (in the decidua).

#### In Vitro Decidualization of Primary HESCs

The expression of *ADGRG2* and *ADGRL4* was quantified and compared in control and *in vitro* decidualized conditions (day 6) using qPCR. A significant increase in expression was seen for *ADGRL4* following six days of *in vitro* decidualization in both HESC lines, thus confirming that its expression is altered with decidualization (Figure12B) Unlike ADGRL4, a significant increase in *ADGRG2* mRNA expression was not observed (Figure 11A).

**Figure 11.**
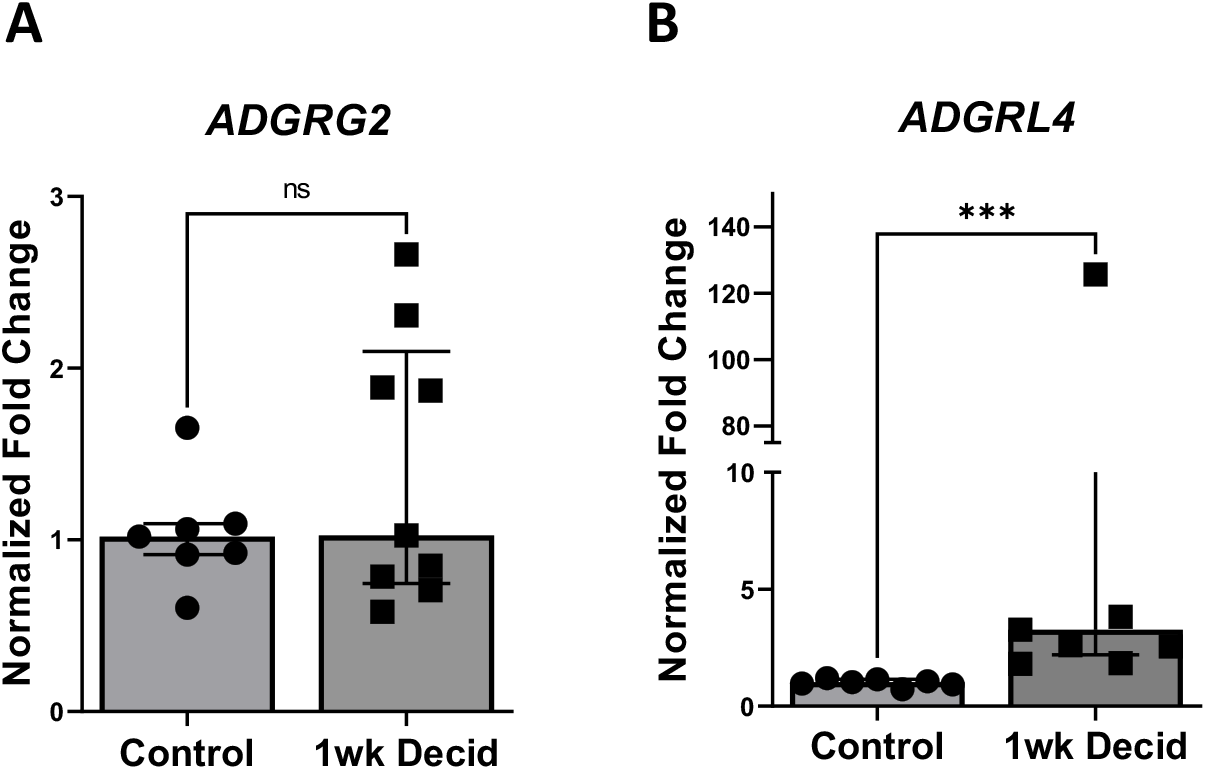
qPCR analysis of (A) *ADGRG2* and (B) *ADGRL4* mRNA expression in primary human endometrial stromal cells that had undergone one week of in vitro decidualization.

## DISCUSSION

GPCRs are major regulators of the reproductive-axis and are targeted pharmacologically in the treatment of infertility. Studies in mouse models have revealed critical roles for GPCRs in the regulation of early pregnancy at the level of endometrial receptivity, embryo implantation and decidualization (Yoo et al. 2017). Less is known about the roles of GPCRs in regulating these events in women, but among the little that is known, an in vitro study showed a potential role for ADGRG2 in regulating decidualization (Yoo et al. 2017). ADGRs represent the second largest GPCR subfamily in humans and are represented by least 32 coding *ADGR* genes. To better understand the roles of ADGRs in regulating the development of the secretory endometrium in women, we began by analyzing their expression in normo-ovulatory women across the menstrual cycle using a published dataset (Talbi et al. 2006). We also studied their expression in women in natural cycles and in OS cycles in the PO period and MS phase of the secretory phase from a prospective cohort.

Our study revealed that members of the *ADGR* gene family are dynamically expressed across the natural menstrual cycle and early mouse pregnancy. Their expression also appears to be altered by ovarian stimulation, during which supraphysiologic levels of E2 and an early rise in P4 are seen (Kalakota et al. 2022). The differential gene expression observed between the proliferative and secretory phase of the menstrual cycle, along with changes in expression seen with OS suggest that *ADGR* expression is hormonally regulated by E2 and P4. Both E2 and P4 are also responsible for the transition of endometrium into a secretory phenotype following ovulation, while P4 and cAMP give rise to the predecidua which regulates embryo implantation. Thus, the changes in *ADGR* expression seen across the menstrual cycle and in early mouse pregnancy suggest roles in the development of an endometrium that is receptive to embryo implantation.

The role of *ADGRG2* in particular has been investigated in both human and mouse decidualization and is used clinically as a marker of endometrial receptivity on the endometrial receptivity array (ERA) (Diaz-Gimeno et al. 2011). The ERA is a molecular assay comprised of 238 genes and is utilized to determine a women’s personal window of implantation (Diaz-Gimeno et al. 2011). The genes selected for use on the ERA were examined across the early secretory phase, anywhere from 1 to 5 days following the surge in luteinizing hormone, and mid secretory phase (LH+7). Those genes that demonstrated an absolute fold change >3 and FDR <0.05 were selected for inclusion. Among these genes is *ADGRG2* which exhibited a 5.5-fold decrease in expression between the early to mid-secretory phase (Diaz-Gimeno et al. 2011). Conversely, another study in human endometrial stromal cells (HESCs) demonstrated increased *ADGRG2* expression with decidualization and furthermore identified a functional role as decidualization in HESCs was impaired after knockdown of *ADGRG2* (Yoo et al. 2017). In our investigation, no significant changes were seen across the menstrual cycle or with ovarian stimulation for *ADGRG2* in the in-silico and prospective RNA seq analyses. However, when expression was tested in additional endometrial biopsies, a significant increase was seen from the proliferative to the mid secretory phase. In vitro studies also demonstrated an increasing trend in *ADGRG2* expression after in vitro decidualization of primary HESCs, however the change in expression was not noted to be significant.

*ADGRG2* has also been identified as a key regulator of decidualization in mouse pregnancy and previous studies have demonstrated significant increases in mRNA and protein expression from E0.5 to E7.5 (Yoo et al. 2017). Our data also demonstrated expression of *ADGRG2* at E7.5 but a significant increase in expression was only seen between E5.5 and E6.5 rather than between E0.5 and a later time point. In addition to ADGRG2, ADGRD1 has also been implicated in mouse embryo transit from the oviduct but was not found to be of significance with respect to mouse decidualization in this study.

G protein coupled receptors are ubiquitous proteins involved in almost every biological processes in humans, including acquisition of endometrial receptivity (Diaz-Gimeno et al. 2011). For this reason, it is imperative to better understand their role in the endometrium and their alterations in expression under ovarian stimulation. Despite doubling the number of IVF cycles performed annually over the past decade, the live birth rate per cycle across all ages, has only increased by a modest 5% since 2010.^3^ OS is associated with reduced implantation rates (Horcajadas, Pellicer, and Simon 2007), and there remains a need for novel and effective therapies to address this problem. Investigating the role of ADGRs in female reproduction could advance the field of assisted reproduction by possibly uncovering new genes that are involved in endometrial receptivity, thereby expanding the panel of genes on the ERA. Additionally, ADGRs could offer targets for novel prophylactic or corrective interventions to improve impaired endometrial receptivity following IVF.

## ABBREVIATIONS

(OS): Ovarian stimulation
(E2): Human endometrial stromal cell Estradiol
(P4): Progesterone
(ADGR): Adhesion G protein-coupled receptors
(EMB): Endometrial biopsies
(NC): Natural cycle
(PO): Peri-ovulatory
(MS): Mid-secretory
(FDR): False detection rate
(DEGs): Differentially expressed genes
(HESC): Human endometrial stromal cell
(LBR): Live birth rate
(LH): Luteinizing hormone
(EMB: Endometrial biopsy
(GPCRs): G protein-coupled receptors
(GnRH): Gonadotropin releasing hormone
(FSH): Follicle stimulating hormone
(HG): High-density human genome
(ESTs: Expressed sequence tags
(hCG): human chorionic gonadotropin
(ECLIA): Electrochemiluminescence immunoassay
(H&E): Hematoxylin and eosin
(IQR): Interquartile range
(PCOS): Polycystic ovary syndrome
(BMI): Body mass index
(SHBG): Sex hormone binding globulin
(AMH): Anti-Müllerian hormone
(ERA): Endometrial receptivity array
(PCA): Principal component analysis

## ACKNOWLEDGEMENTS

Robert Dubin, PhD, Xusheng Zhang, MS and Shahina Maqbool, PhD from the Computational Genomics Core at Albert Einstein College of Medicine for their contributions in the bulk RNA sequencing and primary analysis.

Anat Chemerinski, MD for her contributions in patient recruitment.

Simona Alomary, MD for her contributions with conducting experiments.

## AUTHOR CONTRIBUTIONS

N.R.K., N.C.D, and A.V.B designed the study, performed data analysis, and critically reviewed the manuscript. N.R.K, T.W., Q.Z and L.G. conducted the experiments. S.S.M provided critical feedback with experimental planning. N.R.K wrote the initial draft of the manuscript. A.L. performed the bioinformatic analyses, wrote and revised the manuscript. N.R.K., N.C.D and A.V.B prepared the final draft of the manuscript.

## DATA AVAILABILITY

All data generated or analyzed during this study are included in this article. The prospective human data set has been submitted to GEO under GSE220044.

## DECLARATION OF COMPETING INTERESTS

The authors declare that there are no competing financial, personal, or professional competing interests.

## Notes

https://www.ncbi.nlm.nih.gov/geo/query/acc.cgi?acc=GSE220044

### Competing Interest Statement

The authors have declared no competing interest.

### Summary of Updates

Removal of figure 5 as well as edits to supplemental table 1, 3, and 4.

