## Supplemental Tables for "Endometrial adhesion G protein-coupled receptors are dynamically expressed across the menstrual cycle and expression is altered by ovarian stimulation"

**Supplemental Table 1.** Demographic data of subjects selected from Dataset Record GDS2052 (Endometrium throughout the menstrual cycle) for an in-silico analysis**.**

| Sample ID | Age | Biopsy Type | Hysterectomy Type | Cycle Stage | Pathology |
|---|---|---|---|---|---|
| GSM109814 | 33 | Hysterectomy | Abdominal | Proliferative | Leiomyoma |
| GSM109815 | 31 | EMB | - | Proliferative | - |
| GSM109816 | 32 | EMB | - | Proliferative | - |
| GSM109817 | 34 | EMB | - | Proliferative | - |
| GSM109820 | 46 | Hysterectomy | Supracervical | Early Secretory | Leiomyoma |
| GSM109821 | 48 | Hysterectomy | Abdominal | Early Secretory | Leiomyoma |
| GSM109822 | 44 | Hysterectomy | Vaginal | Early Secretory | Pelvic Organ Prolapse, Adenomyosis |
| GSM109824 | 49 | Hysterectomy | Laparoscopic Assisted Vaginal | Mid Secretory | Leiomyoma (Without Endometriosis) |
| GSM109825 | 42 | Hysterectomy | Vaginal/Abdominal | Mid Secretory | Pelvic pain (Without Endometriosis) |
| GSM109826 | 42 | Hysterectomy | Vaginal | Mid Secretory | Pelvic Organ Prolapse |
| GSM109827 | 46 | Hysterectomy | Vaginal | Mid Secretory | Pelvic Organ Prolapse, Stress Urinary Incontinence |
| GSM109828 | 30 | EMB | - | Mid Secretory | - |
| GSM109829 | 34 | EMB | - | Mid Secretory | - |
| GSM109830 | 23 | EMB | - | Mid Secretory | - |
| GSM109831 | 33 | EMB | - | Mid Secretory | - |
| GSM109834 | 43 | Hysterectomy | Abdominal | Late Secretory | Leiomyoma, Adenomyosis |
| GSM109835 | 39 | Hysterectomy | Laparoscopic Assisted Vaginal | Late Secretory | Leiomyoma |
| GSM109836 | 39 | Hysterectomy | Abdominal | Late Secretory | Leiomyoma |
| GSM109837 | 41 | EMB | - | Late Secretory | Right Ovarian Cyst |
| GSM109838 | 43 | Hysterectomy | Vaginal | Late Secretory | Pelvic Organ Prolapse |
| GSM109839 | 44 | Hysterectomy | Abdominal | Late Secretory | Leiomyoma |

**Supplemental Table 2**. List of the *ADGR* genes uncovered in the GDS2052 dataset and their corresponding probes in the Affymetrix high-density human genome (HG) U133 Plus 2.0 Array.

| Gene | Probe |
|---|---|
| *ADGRA1* | 239221_at |
| *ADGRA2* | 65718_at<br>221814_at |
| *ADGRA3* | 1569542_at<br>210473_s_at |
| *ADGRB1* | 206083_at |
| *ADGRB2* | 204966_at |
| *ADGRB3* | 205638_at<br>211568_at |
| *ADGRD1* | 232267_at |
| *ADGRD2* | 216289_at |
| *ADGRE1* | 207111_at |
| *ADGRE2* | 232009_at<br>207610_s_at |
| *ADGRE3* | 210724_at |
| *ADGRE5* | 202910_s_at |
| *ADGRF1* | 220907_at<br>235988_at<br>238689_at<br>236489_at |
| *ADGRF3* | 236608_at |
| *ADGRF4* | 237690_at |
| *ADGRF5* | 212950_at<br>212951_at |
| *ADGRG1* | 206582_s_at<br>212070_at |
| *ADGRG2* | 212070_at<br>206002_at |
| *ADGRG3* | 220404_at<br>1553723_at |
| *ADGRG5* | 229971_at |
| *ADGRG6* | 233887_at |
| *ADGRG7* | 1553296_at |
| *ADGRL1* | 203488_at<br>219145_at<br>47560_at |
| *ADGRL2* | 1569850_at<br>206953_s_at |

| | |
|---|---|
| *ADGRL3* | 209867_s_at<br>209866_s_at<br>236264_at |
| *ADGRL4* | 219134_at |
| *ADGRV1* | 215396_at<br>223582_at<br>224275_at<br>240631_at |

**Supplemental Table 3**. Demographic data of subjects recruited in the proliferative, peri-ovulatory and mid secretory phase of natural cycles.

| Subject | Cycle Stage | Age | Estradiol Level at Biopsy (pg/ml) | Progesterone Level at Biopsy (ng/ml) |
|---|---|---|---|---|
| **1** | PROF (CD 10) | 28 | 176 | 0.3 |
| **2** | PROF (CD 12) | 26 | 249 | 0.8 |
| **3** | PROF (CD 11) | 27 | 272 | 0.2 |
| **4** | PROF (CD 12) | 29 | 82 | 0.2 |
| **5** | NC- PO | 21 | 213 | 2.2 |
| **6** | NC- PO | 31 | 314 | 0.2 |
| **7** | NC- PO | 33 | 105 | 1.4 |
| **8** | NC- PO | 27 | 182 | 0.1 |
| **9** | NC- PO | 28 | 168 | 0.4 |
| **10** | NC- PO | 32 | 158 | 0.5 |
| **11** | NC- MS | 36 | 86.6 | 15.6 |
| **12** | NC- MS | 26 | 156 | 13.2 |
| **13** | NC- MS | 27 | 121 | 4.8 |
| **14** | NC- MS | 28 | 220 | 6.5 |
| **15** | NC- MS | 29 | 442 | 5.7 |

**Supplemental Table 4**. Demographic data of subjects recruited in the peri-ovulatory and mid-secretory phase of ovarian stimulation cycles.

| Subject | Cycle Stage | Age | Number of Oocytes Retrieved | Estradiol Level at Biopsy (pg/ml) | Progesterone Level at Biopsy (ng/ml) | Peak Estradiol (pg/ml) | Peak Progesterone (ng/ml) |
|---|---|---|---|---|---|---|---|
| **1** | OS-PO | 35 | 19 | 176 | 0.3 | 664 | 12.5 |
| **2** | OS-PO | 32 | 32 | 249 | 0.8 | 1047 | 23.7 |
| **3** | OS-PO | 31 | 19 | 272 | 0.2 | 1182 | 13.7 |
| **4** | OS-PO | 33 | 9 | 82 | 0.2 | 800 | 3.5 |
| **5** | OS-PO | 36 | 15 | 213 | 2.2 | 453 | 13.5 |
| **6** | OS-MS | 39 | 9 | 314 | 0.2 | 585 | 25.4 |
| **7** | OS-MS | 42 | 12 | 105 | 1.4 | 497 | 11.7 |
| **8** | OS-MS | 32 | 9 | 182 | 0.1 | 289 | 4.5 |
| **9** | OS-MS | 36 | 11 | 168 | 0.4 | 1082 | 29.8 |

**Supplemental Table 5**. Demographic data of subjects recruited in the peri-ovulatory and mid-secretory phase of ovarian stimulation cycles. Values listed as medians with interquartile range.

| Cycle Stage | Age | Estradiol Level at Biopsy (pg/ml) | Progesterone Level at Biopsy (ng/ml) |
|---|---|---|---|
| PROF | 27.5 ± 2.5 (26.2-28.8) | 212.5±160.8 (105.5-266.3) | 0.3±0.5 (0.2-0.7) |
| NC-PO | 29.5±6.8 (25.5-32.3) | 175.0±93.5 (144.8-238.3) | 0.5±1.4 (0.2-1.6) |
| OS-PO | 33.0± 4.0 (31.5-35.5) | 213.0±131.5 (129.0-260.5) | 0.3±1.3 (0.2-1.5) |
| NC-MS | 28.0±6.0 (26.5-32.5) | 156.0±227.2 (103.8-331.0) | 6.5±9.1 (5.3-14.4) |
| OS-MS | 37.5±8.3 (33-41.3) | 175±160.2 (120.8-281.0) | 0.3±1.1 (0.1-1.2) |

**Supplemental Table 6**. Human primer sequences.

| Gene | Forward Primer 5'-3' | Reverse Primer 5'-3' |
|---|---|---|
| ***ADGRB2*** | GTGGAGGACTTCATTCACCTGG | GGAACGTGATGTCACTGGACAC |
| ***ADGRF1*** | ACGGCTCTTTCAGAGTGTTCGG | CAGAGGACTCACACTTGGCTGT |
| ***ADGRG2*** | TGAAAGGGGTTGAATCTCCC | CTGCAGGATCCCATTGTCTG |
| ***ADGRL1*** | GGAGGAGTACCCGAGAAAGA | CCAGGTTGTTGTAGAGGATGAA |
| ***18SN5*** | CCCGTTGAACCCCATTCGTGA | GCCTCACTAAACCATCCAATCGG |

**Supplemental Table 7**. Mouse primer sequences.

| Gene | Forward Primer 5'-3' | Reverse Primer 5'-3' |
|---|---|---|
| ***Adgra1*** | GCTCTGACCTTCACCGTGTTTG | GTTCCTAGCAGTCACACCGATC |
| ***Adgra2*** | CGTGGTCTATGTGGCTCAGATG | AGCCACAGAAGGTGCTCATCCA |
| ***Adgra3*** | GTGGTGTTTGTGGGAGGAATA | CCGTGGCAAGGGTAGAATAAT |
| ***Adgrb1*** | CTCCACCATTGATGTCCTGAGG | TCTTCTGCCAGCAGGTTGCTGA |
| ***Adgrb2*** | CTCTGTGGACATCCTAAGGAACG | CCATGAAGCTCACCACCTGGAA |
| ***Adgrb3*** | CGCCATTTTGGCTCAGCAACCT | CCTCCATAGTGCTGCGTACACA |
| ***Adgre1*** | CGTGTTGTTGGTGGCACTGTGA | CCACATCAGTGTTCCAGGAGAC |
| ***Adgre4*** | GCTCTCCATCTGCCTTTTCCTG | GGAAGCCAAGTAGAGGTAGTGC |
| ***Adgre5*** | CTGTCTTCCTGACCAACACGAAC | GTTGTGGGCTTTCCAGAAGGCA |
| ***Adgrf1*** | CCTCCAGAACTCCTCTTTGCCA | CCAAAGGAGATGCTGTTACACGG |
| ***Adgrf2*** | AGGAGAAGGTGACATCCAGACC | ATGTTGCCCTCACTGTCTGGAG |
| ***Adgrf3*** | GGAAATGCTGCACACTGTGACC | GAATCTTGTCTGTGGTCGTCAGG |
| ***Adgrf4*** | CAGATTGTCCCTGGTAGGTTTC | CCCTACACGTCAGGTGATTTAC |
| ***Adgrf5*** | GGAAGAACAGGACATCCGCTCA | CCAGGAGTTCAAGGCAGACTTG |
| ***Adgrg1*** | TCCAGGCATACTCGCTGTTGCT | CTTCTCACCCAGGACTTGGCTA |
| ***Adgrg2*** | CCCTTCCTCACCAGAAGAG | ATAAGGGCATGATCAAGGGG |
| ***Adgrg3*** | GCACAGACTCTCACTCGCATCT | GAAACAGGCTGATGCTCAGAGC |
| ***Adgrg4*** | GAAGGAAAGTGTTCGGGAGCAG | GTTGTTGACATGAGCGATGGTGT |
| ***Adgrg5*** | GCCTACATTCGCCGATACTTGC | CCTGGCTTTTGGAGAGTGAGGT |
| ***Adgrg6*** | AGAGGATGGACTGAGGCTGTGT | CCAGGCTTGTTTGGACATGGTTG |
| ***Adgrg7*** | TGCAGCCACACGACCAATTTCG | GGATAAGGCACAGCCGATGTTG |
| ***Adgrl1*** | GTGGTGGAGACAGTGGATAAC | TAGAAGCATGGTGGCTGTATG |
| ***Adgrl2*** | AGGAGCTGAAGCCGAGTGAGAA | CCAGGATTCCAAAGCCTCAGCT |
| ***Adgrl3*** | CTCTTGCAGAGCCTATGTCCAG | CAGTGTCAAGCAACATGGTGGC |
| ***Adgrl4*** | TCCAAAGCACCAGGACCACGAT | GCCAGCAATGATAGAGCAGACC |
| ***18S*** | GTAACCCGTTGAACCCCATT | CCATCCAATCGGTAGTAGCG |

**Supplemental Table 8**. Differentially expressed *ADGR* genes between the proliferative vs early, mid and late secretory phase in the GDS2052 dataset and their corresponding probes in the Affymetrix high-density human genome (HG) U133 Plus 2.0 Array.

| Gene | Probe | Proliferative | Early Secretory | P-adjusted |
|---|---|---|---|---|
| ADGRE5 | 202910_s_at | 191 | 113 | 0.032 |
| **Gene** | **Probe** | **Proliferative** | **Mid Secretory** | **P-adjusted** |
| ADGRB2 | 204966_at | 37 | 25 | 0.006 |
| ADGRF1 | 235988_at | 37 | 110 | 0.009 |
| ADGRF1 | 238689_at | 31 | 130 | <0.001 |
| ADGRF1 | 236489_at | 19 | 70 | <0.001 |
| ADGRL1 | 203488_at | 267 | 150 | <0.001 |
| ADGRL1 | 219145_at | 149 | 101 | <0.001 |
| ADGRL1 | 47560_at | 288 | 170 | 0.002 |
| **Gene** | **Probe** | **Proliferative** | **Late Secretory** | **P-adjusted** |
| ADGRA2 | 65718_at | 369 | 172 | <0.001 |
| ADGRA2 | 221814_at | 899 | 372 | <0.001 |
| ADGRB2 | 204966_at | 37 | 27 | 0.033 |
| ADGRB3 | 205638_at | 28 | 17 | 0.002 |
| ADGRE2 | 232009_at | 20 | 26 | 0.016 |
| ADGRE2 | 207610_s_at | 27 | 52 | 0.015 |
| ADGRF1 | 235988_at | 37 | 81 | 0.033 |
| ADGRF1 | 238689_at | 31 | 141 | <0.001 |
| ADGRF1 | 236489_at | 19 | 47 | 0.009 |
| ADGRF5 | 212950_at | 142 | 218 | 0.043 |
| ADGRF5 | 212951_at | 155 | 187 | 0.031 |
| ADGRG3 | 220404_at | 62 | 114 | <0.001 |
| ADGRG3 | 1553723_at | 31 | 71 | <0.001 |
| ADGRG7 | 1553296_at | 18 | 21 | 0.044 |
| ADGRL1 | 203488_at | 267 | 141 | <0.001 |
| ADGRL1 | 219145_at | 149 | 80 | <0.001 |
| ADGRL1 | 47560_at | 288 | 128 | <0.001 |
| ADGRL3 | 209867_s_at | 119 | 74 | 0.022 |
| ADGRL4 | 219134_at | 70 | 145 | 0.044 |

Expression values are mean normalized hit counts, and differential gene expression group cross-comparisons were performed using DESeq2 (v1.26.0). Significant differentially expressed gene thresholds were set at FDR adjusted p <0.05.

**Supplemental Table 9**. Differentially expressed *ADGR* genes between the early vs mid and late secretory phase in the GDS2052 dataset and their corresponding probes in the Affymetrix high-density human genome (HG) U133 Plus 2.0 Array.

| **Gene** | **Probe** | **Early Secretory** | **Mid Secretory** | **P-adjusted** |
|---|---|---|---|---|
| ADGRF1 | 235988_at | 29 | 130 | 0.001 |
| ADGRF1 | 235988_at | 34 | 110 | 0.023 |
| ADGRF1 | 238689_at | 19 | 70 | 0.005 |
| ADGRL2 | 206953_s_at | 126 | 351 | 0.010 |
| ADGRL4 | 219134_at | 49 | 108 | 0.020 |
| **Gene** | **Probe** | **Early Secretory** | **Late Secretory** | **P-adjusted** |
| ADGRA2 | 221814_at | 714 | 372 | 0.002 |
| ADGRA2 | 65718_at | 334 | 172 | 0.001 |
| ADGRD1 | 232267_at | 349 | 154 | 0.022 |
| ADGRE2 | 232009_at | 21 | 26 | 0.004 |
| ADGRE2 | 207610_s_at | 22 | 52 | 0.006 |
| ADGRE5 | 202910_s_at | 113 | 238 | <0.001 |
| ADGRF1 | 238689_at | 29 | 141 | <0.001 |
| ADGRF1 | 235988_at | 34 | 81 | 0.037 |
| ADGRF1 | 236489_at | 19 | 47 | 0.023 |
| ADGRF4 | 1553031_at | 8 | 11 | 0.004 |
| ADGRF5 | 212950_at | 132 | 218 | 0.014 |
| ADGRG1 | 212070_at | 416 | 944 | 0.042 |
| ADGRG3 | 1553723_at | 26 | 71 | <0.001 |
| ADGRG3 | 220404_at | 53 | 114 | <0.001 |
| ADGRL1 | 219145_at | 118 | 78 | 0.001 |
| ADGRL1 | 47560_at | 220 | 128 | 0.003 |
| ADGRL2 | 206953_s_at | 126 | 344 | 0.005 |
| ADGRL4 | 219134_at | 49 | 145 | <0.001 |

Expression values are mean normalized hit counts, and differential gene expression group cross-comparisons were performed using DESeq2 (v1.26.0). Significant differentially expressed gene thresholds were set at FDR adjusted $p < 0.05$.

**Supplemental Table 10**. Differentially expressed *ADGR* genes between the peri-ovulatory and mid-secretory phase of natural and ovarian-stimulated cycles.

| Gene | Peri-ovulatory Natural cycle | Mid-secretory Natural cycle | P-adjusted |
|---|---|---|---|
| *ADGRF1* | 27 | 4229 | <0.001 |
| *ADGRL2* | 4631 | 5494 | 0.501 |
| *ADGRL4* | 1085 | 1617 | 0.014 |
| **Gene** | **Peri-ovulatory Ovarian-stimulated cycle** | **Mid-secretory Ovarian-stimulated cycle** | **P-adjusted** |
| *ADGRF1* | 3 | 2100 | <0.001 |
| *ADGRL2* | 2015 | 8365 | <0.001 |
| *ADGRL4* | 827 | 1749 | <0.001 |
| *ADGRC1* | 1906 | 4825 | 0.012 |
| ADGRL4 | 349 | 154 | 0.022 |
| **Gene** | **Peri-ovulatory Natural cycle** | **Peri-ovulatory Ovarian-stimulated cycle** | **P-adjusted** |
| *ADGRF1* | 27 | 3 | 0.023 |
| *ADGRL2* | 4631 | 2015 | <0.001 |
| *ADGRL4* | 1085 | 827 | 0.182 |
| ADGRL4 | 349 | 154 | 0.022 |
| **Gene** | **Mid-secretory Natural cycle** | **Mid-secretory Ovarian-stimulated cycle** | **P-adjusted** |
| *ADGRF1* | 4229 | 2100 | 0.771 |
| *ADGRL2* | 5494 | 8365 | 0.319 |
| *ADGRL4* | 1617 | 1749 | 0.882 |

Expression values are mean normalized hit counts, and differential gene expression group cross-comparisons were performed using DESeq2 (v1.26.0). Significant differentially expressed gene thresholds were set at FDR adjusted p <0.05.
